# Dynamic and context-dependent modulation of proprioceptive input in primate primary motor cortex

**DOI:** 10.1101/2025.09.21.677658

**Authors:** Junichiro Yoshida, Satomi Kikuta, Woranan Hasegawa, Shinji Kubota, Kazuhiko Seki

## Abstract

During voluntary movement, somatosensory input is attenuated—sensory gating—which may prevent the CNS from being overwhelmed by predictable afferent feedback not essential for motor control. In the cerebral cortex, sensory gating has been demonstrated in primary (S1) and higher-order sensory areas. The primary motor cortex (M1) is the major cortical output relay to the spinal cord and muscles, and proprioceptive feedback is crucial for shaping motor output, much as in S1. However, whether proprioceptive signals to M1 are attenuated during movement, and if so, why, remains unclear. We recorded somatosensory-evoked potentials (SEPs) in M1 of two monkeys performing a wrist task while electrically stimulating muscle and cutaneous afferents innervating forearm extensors and adjacent skin. Both local field potentials and single-neuron recordings revealed significant suppression of muscle-evoked SEPs during Active movement and Static hold phases, providing direct evidence that proprioceptive input to M1 is generally gated during motor execution, as previously reported for cutaneous input. Yet, amidst this suppression, SEPs in a subset of M1 neurons were preserved during Static hold, especially those evoked by antagonist muscle afferents. Because monkeys had to maintain precise joint angles for stable posture, these results suggest that proprioceptive signals from antagonist muscles—likely reflecting spindle activity in lengthening muscles—escape attenuation to provide information essential for joint angle control. Overall, our findings demonstrate that while proprioceptive input to M1 is broadly suppressed during motor output, specific afferent signals from antagonist muscles are selectively maintained in a context-dependent manner to support posture control.

## Introduction

The primary motor cortex (M1) plays a pivotal role in generating voluntary movements through direct corticospinal projections from corticomotoneuronal neurons to spinal motoneurons (Fetz et al., 1980; Phillips & Porter, 1964; Rathelot & Strick, 2009). However, M1 is not solely a motor output area. Anatomical and physiological studies have demonstrated that M1 also receives substantial somatosensory input from the thalamus and the primary somatosensory cortex (Asanuma et al., 1980; Asanuma & Mackel, 1989; Celesia, 1979; Holsapple et al., 1991; Horne & Tracey, 1979; Stepniewska et al., 1993; Tokuno & Tanji, 1993), and M1 neurons are known to exhibit robust somatosensory responses (Lemon, 1981; Rosén & Asanuma, 1972; Strick & Preston, 1982). These findings suggest that M1 functions as a hub for integrating sensory feedback with motor commands (Fetz et al., 1980; Hatsopoulos & Suminski, 2011).

Given that every voluntary movement generates reafferent somatosensory signals, the presence of rich sensory input to M1 poses a fundamental question: how does the brain prevent task-irrelevant or disruptive sensory information from interfering with motor execution (Tomatsu et al., 2023)? Without appropriate modulation, these self-generated sensory signals could corrupt the fine control of voluntary actions (Miall et al., 1985). This problem necessitates a mechanism that filters incoming sensory information based on behavioral context.

The central nervous system addresses this challenge through a phenomenon known as sensory gating, in which somatosensory signals associated with self-generated movement are selectively suppressed (Blakemore et al., 1998; Cohen & Starr, 1987; Lei et al., 2018; Rushton et al., 1981). Sensory gating has been demonstrated across multiple levels of the neuraxis, including the spinal cord (Confais et al., 2017; Seki et al., 2003; Tomatsu et al., 2023), brainstem (Ghez & Pisa, 1972; Kubota et al., 2024), and sensorimotor cortex (Jiang et al., 1990; Seki & Fetz, 2012).

At the spinal level, for example, both muscle and cutaneous afferent inputs are gated during voluntary movement (Confais et al., 2017; Seki et al., 2003; Tomatsu et al., 2023), which contributes to shaping motor commands by appropriately integrating sensory feedback with descending cortical inputs to spinal motoneurons. In contrast, at the cortical level, sensory gating in M1 has been demonstrated only for cutaneous afferent inputs (Seki & Fetz, 2012). Whether muscle afferent signals are also gated remains unclear, and determining their presence is essential for understanding how different types of sensory inputs are processed in M1. If such gating exists, it may not simply mirror the modulation observed for cutaneous afferents, but instead exhibit qualitatively distinct characteristics. This possibility is supported by the fact that muscle afferents convey critical information about limb position and force (Matthews, 1981; Proske & Gandevia, 2009), and play a central role in coordinating multi-joint movement (Bossom J, 1974; Sainburg et al., 1995). Given their fundamental importance in motor execution, muscle afferent input to M1 may be subject to context-dependent modulation that differs in both form and functional impact from that of cutaneous input. This view is further supported by previous findings, which showed changes in M1 responses to mechanical perturbations applied to the arm depending on behavioral context (Evarts & Fromm, 1977; Evarts & Tanji, 1976; Pruszynski et al., 2011).

To investigate this hypothesis, we electrically stimulated peripheral nerves—either muscle or cutaneous forearm nerves—while monkeys performed voluntary wrist movements, and recorded evoked responses in M1. Among M1 neurons that responded to muscle nerve stimulation, the majority exhibited significant suppression during both Active movement and Static hold. Intriguingly, a subset of neurons showed selective preservation of responses, particularly from antagonist muscle input during the Static hold epoch of the task, which required precise control of joint angle to maintain a stable posture. These results suggest that while the general principle of sensory suppression holds, M1 may also selectively maintain specific sensory signals from muscle afferents depending on motor context.

## Materials and Methods

### Subjects

All animal procedures were approved by the Institutional Animal Care and Use Committee at the National Institute of Neuroscience, National Center of Neurology and Psychiatry, Kodaira, Japan. Two adult male macaque monkeys participated in this study: monkey T (*Macaca mulatta*, 6.4 kg, aged 7 years) and monkey N (*Macaca mulatta*, 7.0 kg, aged 7 years), both obtained from HAMRI Co., Ltd. (Ibaraki, Japan). A standard commercial non-human primate diet (AS; Oriental Yeast Oriental Yeast, Tokyo, Japan) was provided daily (120–150 g/day), supplemented with a variety of vegetables, fruits, and grains. Water was available *ad libitum*.

### Surgical procedures

Surgical procedures were conducted under aseptic conditions. Atropine, midazolam and xylazine were administered preoperatively, and anesthesia was maintained with 2.0–3.0% sevoflurane anesthesia. Ampicillin and prednisolone were given postoperatively. To stimulate cutaneous and muscle afferents, tripolar nerve cuff electrodes were implanted on the superficial (SR) and deep (DR) branches of the radial nerve, respectively (Fig.1A). The SR branch innervates a skin patch on the dorsal, radial aspect of the hand, whereas the DR branch predominantly innervates muscle fibers and afferent fibers from forearm extensor muscles and their tendon organs. An additional cuff electrode was implanted proximally on the radial nerve to record afferent volleys elicited by DR or SR stimulation.

**Figure 1.**
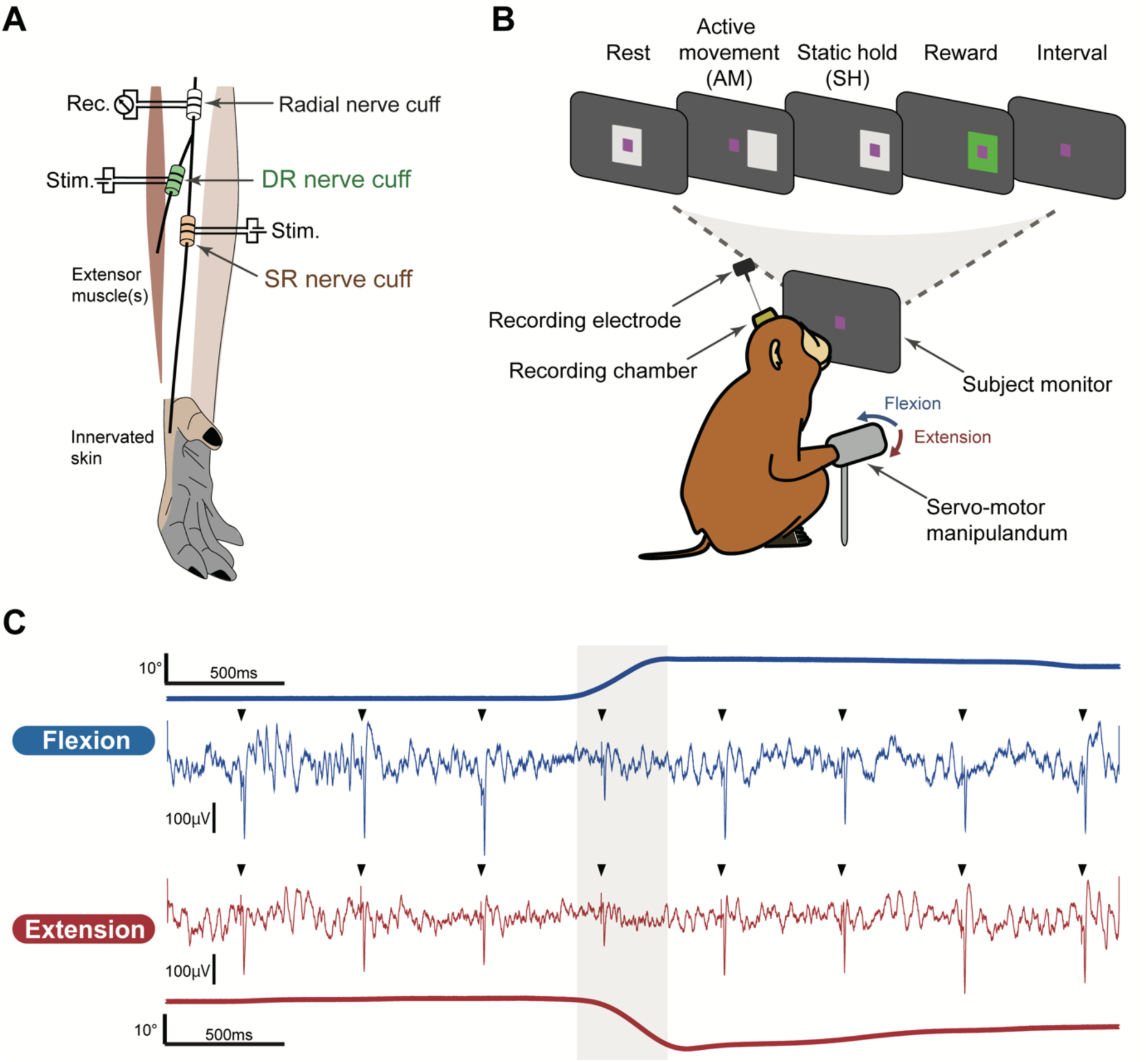
Recording setup. **A,** Peripheral nerve stimulation. Three nerve cuff electrodes were chronically implanted in the superficial radial branch (SR), deep radial branch (DR), and radial stem nerve of the right arm. **B,** Behavioral task. Monkey performed the wrist flexion and extension movement task guided by visual feedback. The task consisted of five epochs: Rest, Active movement, Static hold, Reward, and Interval. **C,** Neural activities and wrist kinematics. A representative example of local field potentials recorded from the primary motor cortex (M1) alongside the angle trajectory (thick line) during a single trial of flexion (blue) and extension (red). The shaded area indicates the joint displacement period. Arrowheads mark the timing of SR nerve stimulation.

In monkey T, pairs of stainless wire electrodes (AS631; Cooner Wire) were implanted into the right forearm muscles for electromyography (EMG) recordings: extensor carpi ulnaris (ECU), extensor carpi radialis (ECR), extensor digitorum communis (EDC), flexor carpi radialis (FCR), flexor carpi ulnaris (FCU), and flexor digitorum superficialis (FDS). Connectors of EMG and nerve cuff electrodes were secured to the skull with dental acrylic anchored by bone screws. A custom-made recording chamber (polyetherimide, hollow square prism, 20 mm per side) was implanted over a craniotomy on the left side of the skull, using dental resin.

### Behavioral task paradigm

Monkeys were trained to perform a wrist flexion and extension movement task, as described previously (Kubota et al., 2024). Briefly, during training and recording sessions the right hand was secured in a cast with fingers extended and the wrist in a neutral position (mid-supination/pronation). The cast was attached to a servomotor-driven manipulandum (Techno-hands Co., Ltd, Yokohama, Kanagawa) to measure wrist torque and angle (Fig.1B). Using the TEMPO system (Version 11.7, Reflective Computing, WA, USA), animals controlled a red cursor on a front-mounted monitor by flexing or extending the wrist. Each trial began with a white center box (Fig.1B, “Rest”). After a 1.0-1.5 s hold, the center box disappeared and a peripheral target appeared, cueing flexion or extension (Fig. 1B, “Active movement”). The cursor had to be maintained within the target for 0.8–1.2 s (Fig.1B, “Static hold”), after which a juice reward was delivered (Fig.1B, “Reward”). The intertrial interval was 1–2 s (Fig.1B, “Interval”). Trials were classified as errors if the cursor was not maintained within the center or the instructed target for the required duration.

### Data recordings

During the recording session, the monkey’s head was restrained with a thermoplastic mask (Uni-frame, Toyo Medic, Tokyo, Japan) (Drucker et al., 2015). A hydraulic micromanipulator (MO-973; NARISHIGE, Japan) was mounted onto the recording chamber via a custom-made X-Y positioning stage. Local field potentials (LFPs) and neuronal spike activity were recorded in the M1 using either stainless steel “Deep array” probes (32 channels at 100 μm or 200 μm spacing, Cambridge Neurotech, Inc., UK) or platinum/iridium S-probes (32 channels at 100 or 200 μm spacing; 64 channels at 100 μm spacing; Plexon Inc., USA). Electrode impedance ranged from 0.1–1.0 MΩ at 1 kHz.

Electrodes were inserted in the precentral cortex with a 0.5 mm grid (Fig. S1A). M1 was identified in three steps. First, we applied low-intensity intracortical microstimulation (0.15-ms biphasic pulses, 333 Hz, trains of 10 pulses) directly to the cortex. The forelimb M1 area was defined by the low thresholds (less than 30 μA) for evoked movement (Fig. S1A; Rathelot & Strick, 2009). Second, electrode locations were verified by computed topography (CT) and magnetic resonance (MR) images (Fig. S1B; Miocinovic et al., 2007). Third, we confirmed the characteristics of M1 LFP waveform, a small positive deflection preceding a negative component (Seki & Fetz, 2012).

The DR or SR nerve was stimulated with biphasic pulses (0.1-ms duration) at 2 Hz. The threshold current that evoked an incoming volley from DR or SR stimulation was recorded at the beginning of the session. In each session, the stimulus intensity was set to 1.5–2.0 times the threshold for DR or SR stimulation. SR- or DR-evoked LFPs and neuronal activity in M1 were recorded while animals performed the task (Fig. 1B).

Signals were amplified (×20) and digitized with the AlphaLab SNR system (Alpha Omega Engineering Ltd., Israel) at 22 kHz (neuronal spikes), 1375 Hz (LFPs), and 2.75 kHz (wrist torque and angle). An Ag–AgCl ball electrode placed on the scar tissue overlying the cortical surface served as the ground/reference. LFPs were low-pass filtered at 200 Hz. Spiking activity was band-pass filtered between 200 and 9000 Hz. Single-unit activity was manually isolated offline using spike-sorting software (Offline Sorter 4.11, Plexon Inc., USA). The accuracy of offline sorting was verified by inter-spike interval histograms and computing autocorrelograms.

### Data analysis

For the epoch-based analysis, we defined three task epochs: Rest, Active movement (AM), and Static hold (SH). The Rest epoch was defined as the period from the appearance of the center box to its disappearance. The Active movement epoch was defined as the period from the disappearance of the center box until the peripheral target was entered. The Static hold epoch was defined as the period from target entry until 0.8–1.2 s later.

We quantified the epoch-dependent modulation of stimulus-evoked LFPs by compiling and averaging across a session consisting of at least 80 trials. The size of LFP evoked by SR and DR stimulation was measured as the peak area and peak amplitude under baseline from response onset to the negative peak of the averaged waveform (Fig. S1B). Onset was defined as the time at which the waveform crossed four standard deviations (SDs) relative to baseline (20–40 ms before stimulation), i.e., the first crossing below -4 SD (onset). Responses with onset latencies longer than 15 ms were excluded from subsequent analysis to avoid contamination by reafferent inputs (e.g., muscle twitches).

Peristimulus time histograms (PSTHs; 0.5-ms bins) were constructed for isolated neuronal action potentials aligned to stimulus pulses. For each neuron, the response probability was computed as spikes per bin divided by the number of stimulation pulses, and summarized in the PSTH. Neurons were classified as responsive or non-responsive to DR or SR stimulation based on the positive peak area in the PSTH (Fig. S1D). Peak onset and offset were defined as crossing of +2 SD relative to baseline firing probability (20–50 ms before stimulation). Classification was performed using pulses delivered during the Rest epoch (i.e., outside movement period) to avoid underestimating responsiveness due to movement-related suppression. Neuronal responses with onset latencies longer than 15 ms were also excluded. For epoch-dependent modulation of neuronal responsiveness, we evaluated the PSTH positive peak area within each task epoch. To ensure unbiased estimates, only epochs with more than 10 stimulation pulses were analyzed.

### Quantification and statistical analysis

Unpaired t-test was used to compare onset latency and peak amplitude between groups (Figs. 2B, C and 6). A two-way ANOVA followed by Tukey’s *post hoc* test assessed epoch-dependent modulation of stimulus-evoked LFPs or spike activity across task epochs (Figs. 3B, 4B and D). A binomial test was used to compare the PSTH peak areas between flexion and extension trials (Figs. 5A, B). Finally, Fisher’s exact test compared the proportions of classified neurons between flexion- and extension-suppressed populations (Tables S2, S3).

**Figure 2.**
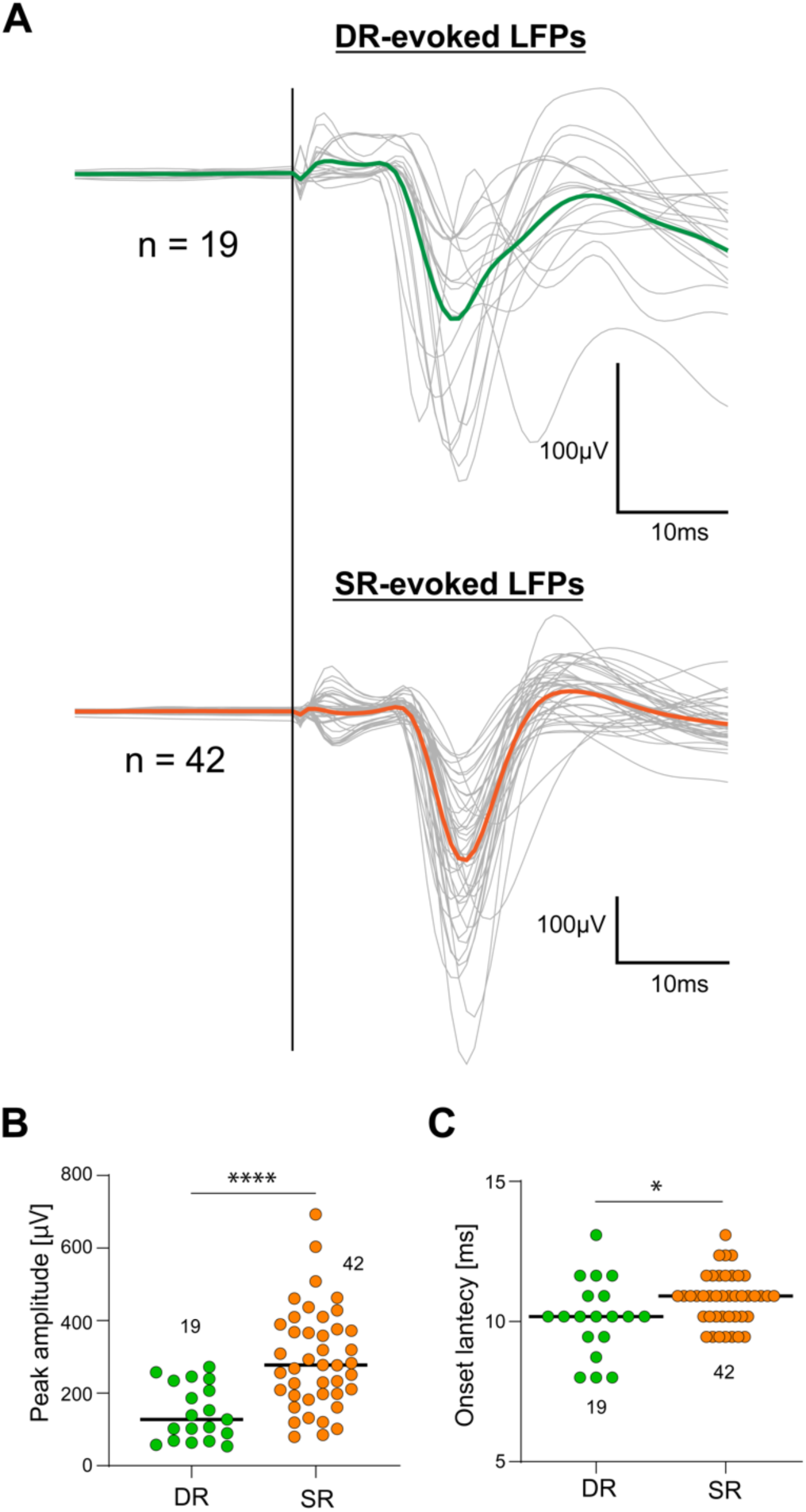
Evoked field potentials in M1 by DR and SR nerve stimulation. **A,** Stimulus-evoked local field potentials (LFPs). DR- and SR-evoked LFPs at each recording site are overlaid in grey, with averaged traces shown in green (DR) and orange (SR). The black vertical line indicates the timing of nerve stimulation. **B, C,** Peak amplitude (B) and onset latency (C). Each dot represents individual data. Numbers above or below the columns indicate the number of data points per group. * p<0.05, **** p<0.0001, unpaired t-test.

**Figure 3.**
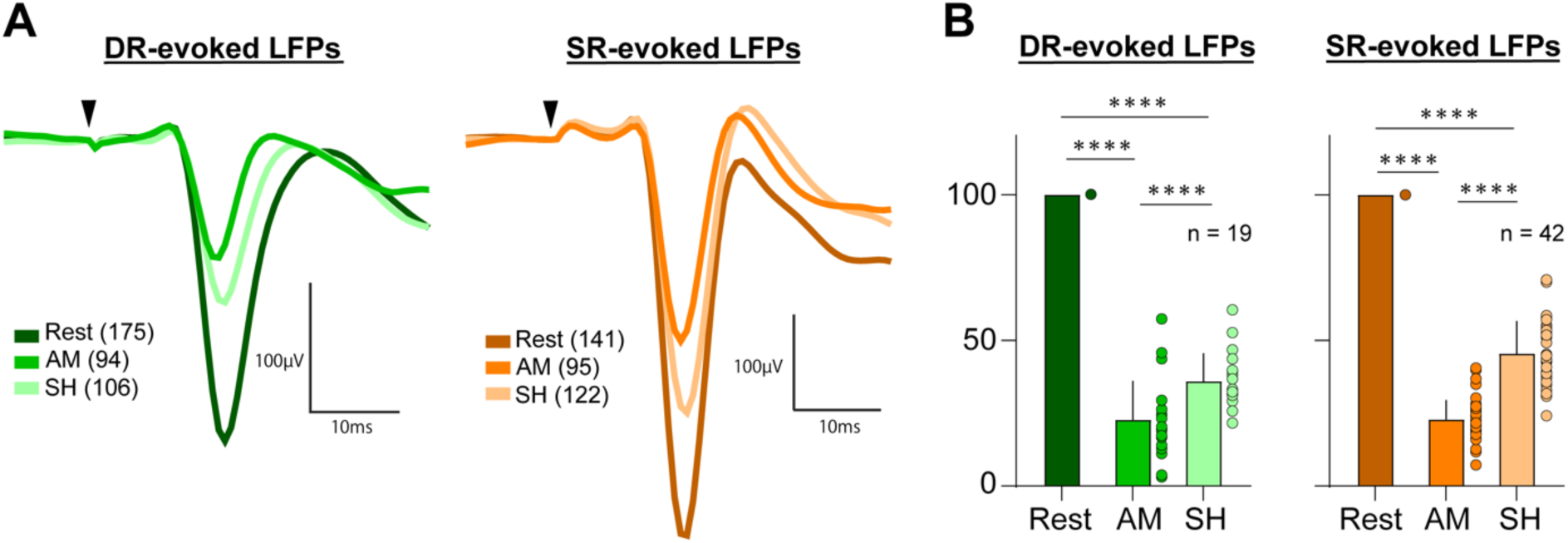
Task-dependent modulation of evoked LFPs in M1. **A,** Representative DR- and SR-evoked LFPs. DR- (left) and SR- (right) evoked LFPs are averaged across trials for the analyzed task epochs: Rest, Active movement (AM) and Static hold (SH). Arrowheads indicate the timing of stimulation delivery, and numbers in parentheses indicate the number of stimulation pulses in a given behavioral epoch. **B,** Peak area quantification. The peak area of DR- (left) and SR- (right) evoked LFPs are shown for the analyzed task epochs. Dots represent individual data. **** p<0.0001, two-way ANOVA with Tukey’s *post hoc* test.

**Figure 4.**
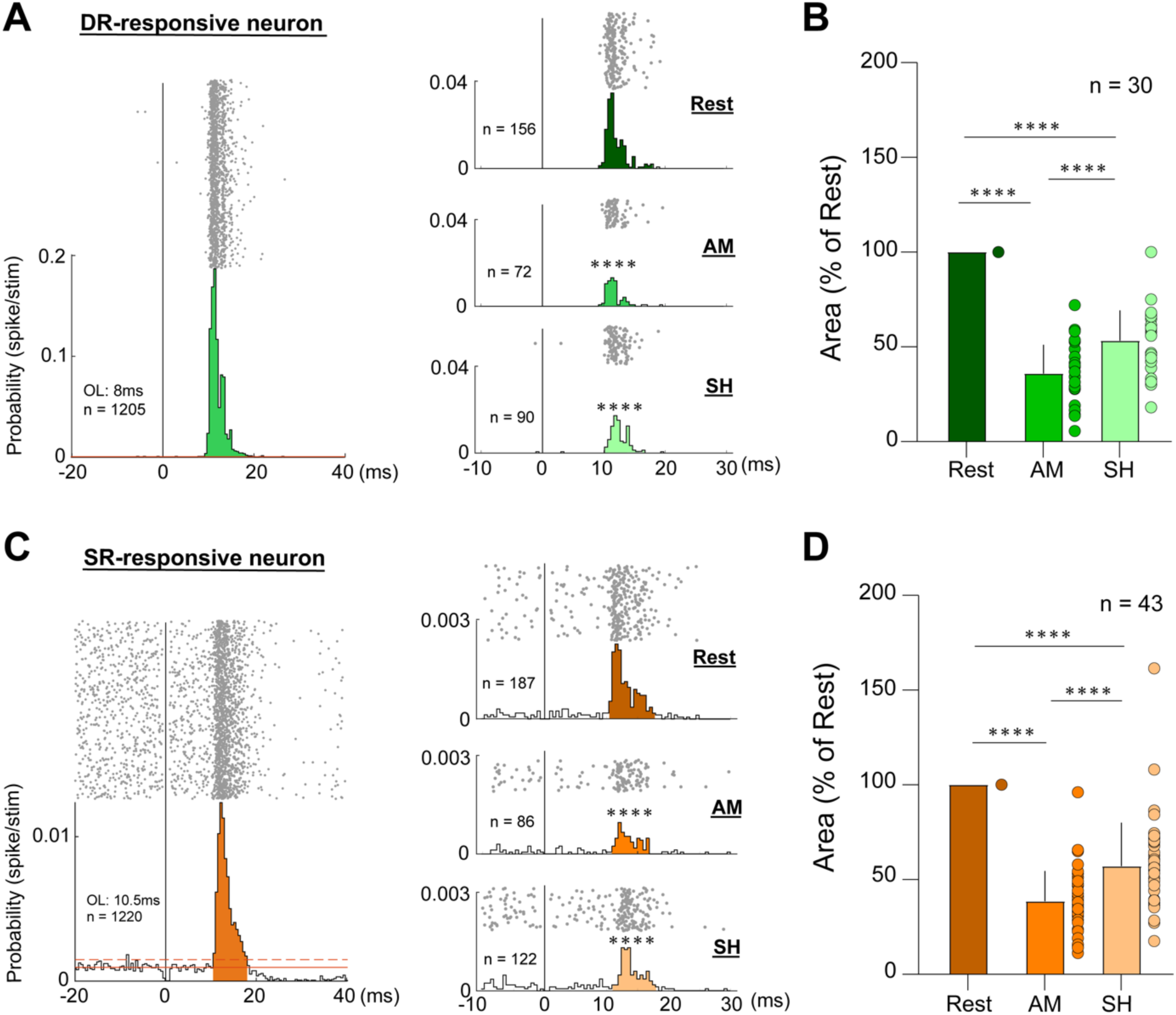
Modulation of DR- and SR-evoked response in M1 neurons during the wrist flexion and extension movement task. **A,** Example of DR-responsive neuron. Raster plot and its peristimulus time histogram (PSTH) aligned to DR stimulation. The left panel shows the response over the task, and the right panel shows the response at the analyzed task epochs: Rest, Active movement (AM) and Static hold (SH). Vertical black line marks the timing of stimulation delivery. Shaded areas indicate PSTH peak, i.e., peak area. PSTH bin size, 0.5 ms; OL, onset latency; n, number of nerve stimulations. **** p<0.0001, binomial test. **B,** Quantification of DR response (mean ± SD). Mean peak area of DR-evoked responses across the task epochs (normalized by the mean peak area at the Rest epoch). Dots represent individual neurons. **** p<0.0001, two-way ANOVA with Tukey’s *post hoc* test. **C and D,** SR-responsive neurons. same as A and B, but for SR stimulation.

**Figure 5.**
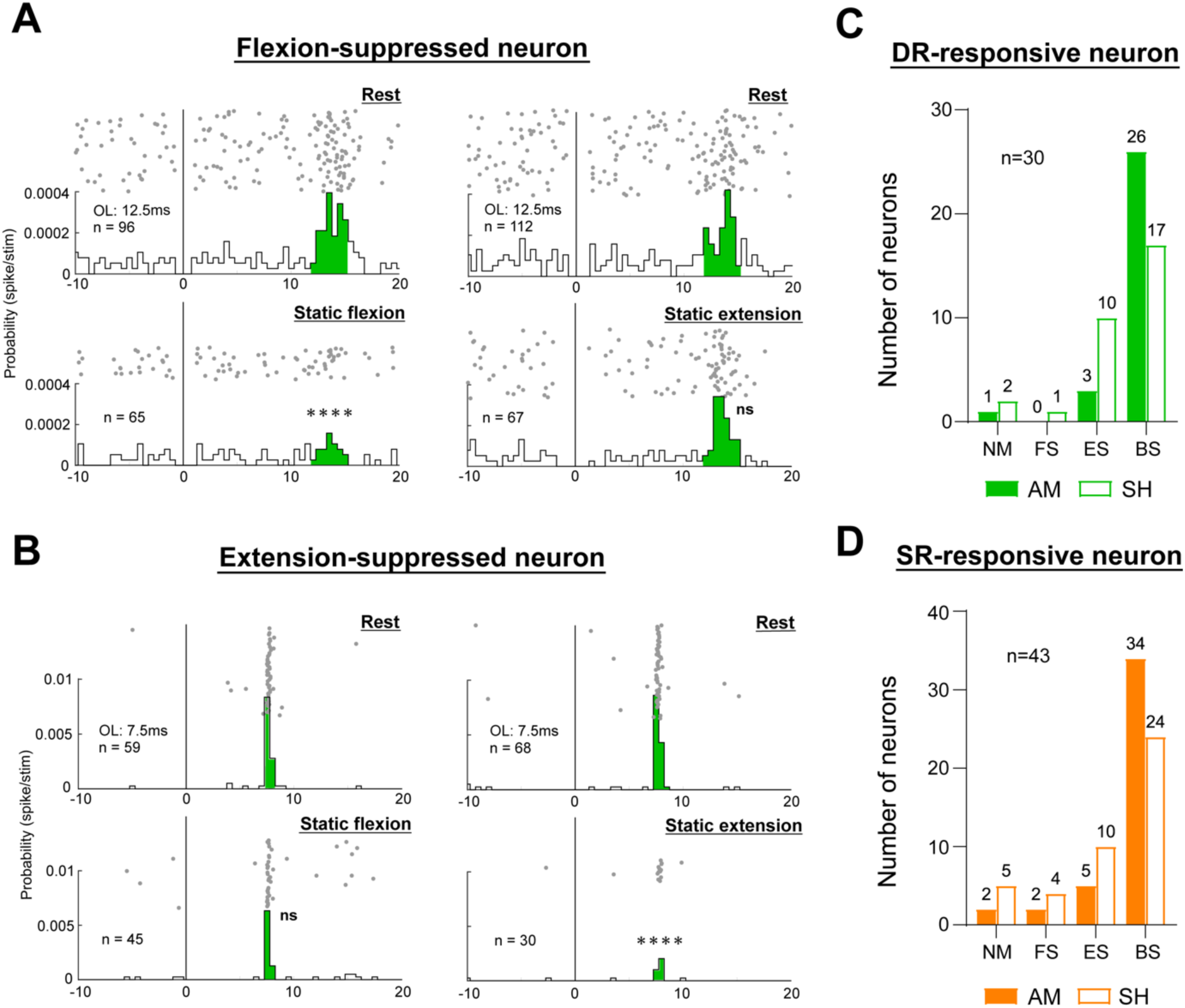
Directional modulation of DR- and SR-responsive neurons during wrist movement. **A,** Example of Flexion-suppressed DR-responsive neuron. Raster plot and PSTH separated by trials (flexion: left, extension: right), and by epoch (Rest: top, Static hold: bottom). Vertical black line marks the timing of stimulation delivery. Shaded areas indicate PSTH peak, i.e., peak area. PSTH bin size, 0.5 ms; OL, onset latency; n, number of nerve stimulations. **** p<0.0001, ns: not significant, binomial test. **B,** Example of an Extension-suppressed DR-responsive neuron. **C and D,** Summary of suppression patterns. Distribution of suppression types in DR- (C) and SR-responsive neurons (D) during the Active movement (AM) and Static hold (SH) epochs. Bars show the number of neurons at each group: NM, non-modulation; FS, flexion-suppressed; ES, extension-suppressed; BS, bilaterally suppressed. Numbers above bars indicate sample size. n, total number of neurons included in this analysis.

Statistical analyses were performed in MATLAB (R2019b, MathWorks Inc., Natick, USA) for the binomial test and in GraphPad Prism 10 (GraphPad Software, Boston, USA) for unpaired t-test, two-way ANOVA with a post hoc test, and Fisher’s exact test.

### Code Accessibility

The codes used in this study are available from the corresponding author upon reasonable request.

## Results

In two monkeys, we made 36 penetrations (20 for monkey T and 16 for monkey N) over M1 (Fig.S1A), either a 32-channel probe (for 33 penetrations) or a 64-channel probe (for 3 penetrations) was inserted. We aimed to record both LFPs and extracellular neuronal spikes simultaneously from each channel in each probe while monkeys performed the wrist flexion and extension movement task. In the following section, we will first describe the response and modulation properties of stimulus-evoked LFPs, followed by those of individual M1 neurons.

### Local field potentials evoked by Muscle afferents in M1

We recorded SR-evoked LFPs in 43 different sites and DR-evoked LFPs in 30 different sites in M1 of two monkeys. Among these, 42 SR- and 19 DR-evoked LFPs exhibited pronounced responses with sufficient signal-to-noise ratio for subsequent analysis (23 SR- and 7 DR-evoked LFPs from monkey T; 19 SR- and 12 DR-evoked LFPs from monkey N).

Fig. 2 shows the waveform of 19 DR-LFPs (A, top), peak amplitude (B) and onset latency (C). The characteristics of SR-LFPs (A, bottom) are also shown for comparison. Although the stimulus intensities applied for DR and SR were comparable, DR-evoked LFPs exhibited an earlier small response compared to SR-evoked LFPs. These differences could be reasonable considering the distributed nature of the cutaneous field potential. In summary, these results clearly showed that muscle afferent signals could reach the macaque M1 under awake, behaving conditions, and affect the excitability of a considerable number of M1 cells that compose the DR-evoked LFPs.

Next, we examined how the sensory inputs from muscle afferents are modulated during a motor task, specifically how the size of DR-evoked LFPs is modulated and whether it differs from the pattern of modulation of SR-evoked LFPs (Seki & Fetz, 2012). Here, we compared two movement epochs, namely Active movement and Static hold epochs, which require ballistic and fine regulation of motor output, respectively. We found that DR-evoked LFPs were significantly suppressed during both Active movement and Static hold epochs. As shown in the representative results (Fig. 3A, left), the size of DR-evoked LFPs was suppressed during both epochs compared to the control. This suppression was more marked in Active movement than Static hold (81% for Active movement and 69% for Static hold), which was comparable to the SR-evoked LFPs (Fig. 3A, right, 73% for Active movement and 55% for Static hold). These characteristics were reproduced in the population analysis (Fig. 3B). For DR-evoked LFPs (n=19), the size decreased by about 77.2±13.7% in the Active movement epoch (p < 0.0001) and 64.0±10.0% in the Static hold epoch (p < 0.0001). For SR-evoked LFPs (n = 30), it decreased by about 77.2±6.8% in the Active movement epoch (p < 0.0001) and 54.6±11.4% in the Static hold epoch (p < 0.0001). These results indicate that, similar to cutaneous afferent input (Seki & Fetz, 2012), muscle afferent input to M1 is also significantly suppressed during both ballistic and static movements. Although previous studies have reported modulation of somatosensory input to macaque M1 during movement (Evarts & Tanji, 1976; Fetz et al., 1980; Flament & Hore, 1988; Lemon et al., 1976), most of these investigations used passive joint movements or mechanical perturbations that simultaneously activated both cutaneous and proprioceptive receptors. As such, they did not clarify whether muscle afferent input alone is modulated during movement. The present results establish that afferent inputs from muscle receptors are subject to sensory gating during voluntary movement.

### Modulation of individual M1 neurons

LFPs recorded in M1 mainly represent the net presynaptic drive into the recorded area (Einevoll et al., 2013; Logothetis et al., 2001; Scherberger et al., 2005). Therefore, results so far indicate that the net presynaptic drive must be decreased during voluntary movement. Next, we examined whether the suppression of presynaptic drive to M1 is effective enough to suppress the recruitment of individual M1 neurons’ activity. For this purpose, we recorded extracellular action potentials, together with the LFPs, of individual 370 neurons from 35 penetrations over M1 in two monkeys (208 from 20 penetrations for monkey T, 159 from 15 penetrations for monkey N). Of the 370 neurons, 316 were examined in response to both SR and DR stimulation. Twenty-eight neurons were recorded under SR stimulation only, and 26 neurons were recorded under DR stimulation only. Among them, 43 (13%) and 30 (9%) neurons showed a significant response probability in Rest, i.e., short-latency peak in the peristimulus time histograms (PSTHs; see methods for detection of PSTH peak). Among these responsive neurons, only 10 neurons exhibited multimodal responses to both DR and SR stimulation. We termed these neurons the “DR-responsive” and “SR-responsive” neurons and used exclusively for the subsequent analysis. Figs. 4A and D show representative examples of a DR-responsive (Fig. 4A) and SR-responsive neuron (Fig. 4D). Although this is expected from the DR-evoked LFPs (Fig. 2A) and SR-evoked LFPs (Fig. 2A, see also Seki and Fetz, 2012), this result suggests the net presynaptic drive from the ascending pathways originating from both SR and DR to the M1 region effectively recruits a subset of M1 neurons.

Next, we compared the response probability of DR- and SR-responsive neurons across Rest, Active movement, and Static hold epochs. Representative examples of neuronal responses during these epochs are shown in Figs. 4A and C (right), corresponding to the same neuron illustrated in Figs. 4A and C (left). Population averages of peak responses (Figs. 4B, D) revealed a significant reduction in response probability during both the Active movement and Static hold epochs compared to Rest. These findings suggest that evoked responses in M1 neurons are suppressed during voluntary movement. This suppression likely reflects modulation—specifically, attenuation—of the movement-related net presynaptic drive to DR- and SR-responsive neurons during movement, as evidenced by the reduced LFP amplitude observed in Fig. 3A. The diminished LFP signals suggest a weakened synaptic input, which in turn may underlie the reduced recruitment of M1 neurons in response to DR and SR stimulation during voluntary movement.

### Task-dependency of sensory attenuation on muscle afferent input

Although the response probabilities of DR- and SR-responsive neurons were reduced during movement, the suppression was far from complete. We observed a 58.9% and 61.7% decrease in response probability for DR- and SR-responsive neurons during Active movement, and 43.8% and 43.3% during Static hold (Figs. 4A and C, right). This partial suppression suggests that the sensitivity of these neurons to afferent input may be at least partially preserved during voluntary movement. If so, it raises an important question: which aspects of afferent input are selectively suppressed, and which are preserved under sensory gating during movement?

To address this question, we examined whether the evoked responses of SR- and DR-responsive neurons differed between wrist flexion and extension movements. The rationale is that SR and DR nerves primarily innervate the wrist extensor muscles and the overlying skin; thus, wrist extension and flexion represent agonistic and antagonistic movements to them, respectively. It is well established that afferent input plays distinct roles during agonistic versus antagonistic movements in the central nervous system—for example, through autogenic and reciprocal reflex pathways in the spinal cord. Based on this, we hypothesized that afferent inputs critical for executing a given movement (e.g., active extension or static hold) might be selectively preserved under sensory gating during voluntary movement.

To address this objective, we compared the suppression profiles of DR-evoked responses during movement between flexion and extension trials. Examples from two DR-responsive neurons are shown in Figs. 5A and B. The neuron in Fig. 5A exhibited minimal attenuation of DR-evoked responses in the extension trials, suggesting that it appeared to receive stronger DR afferent input during the extension Static hold period—corresponding to the agonistic movement. We referred to this type as Flexion-suppressed (FS) neurons. In contrast, the neuron in Fig. 5B showed less attenuation in the flexion trials, indicating greater DR afferent input during the flexion Static hold period—the antagonistic movement. These neurons were termed Extension-suppressed (ES) neurons.

We classified DR-responsive (Fig. 5C) and SR-responsive neurons (Fig. 5D) into four categories: non-modulated (NM), suppressed in both flexion and extension (bilaterally suppressed; BS), and the previously defined flexion-suppressed (FS) and extension-suppressed (ES) types (Table S1).

During Active movement, the response probability of most SR- and DR-responsive neurons was strongly suppressed regardless of movement direction, and these were categorized as BS neurons (Figs. 5C, D; DR: n = 26/30, 87%; SR: n = 34/43, 79%). Notably, only a few neurons fell into the FS and ES types during this epoch (Fig. 5C, D; DR: FS, n = 0/30, 0%; ES, n = 3/ 30, 10%; SR: FS, n = 2/43, 5%; ES, n = 4/43, 9%). Contrastingly, during the Static hold period, a larger proportion of neurons were classified as FS or ES types (Figs. 5C, D; DR: FS, n = 1/30, 3%, ES, n = 10/30, 33%; SR: FS, n = 5/43, 12%, ES, n = 10/43, 23%), although a majority was categorized as the BS group (Figs. 5C, D; DR: n = 17/30, 57%; SR: n = 24/43, 56%). Importantly, within the FS and ES groups, the proportion of ES-type neurons was significantly higher than FS-type neurons specifically among DR-responsive neurons (Table S2; Fisher’s exact test, p < 0.01). This ES-biased distribution was not observed in SR- responsive neurons (Table S3; Fisher’s exact test, p = 0.1433), nor during Active movement in either population (Fisher’s exact test, DR: p=0.2373; SR: p=0.6761).

We further compared the onset latencies between DR-responsive neurons on ES- and BS-types. We found that the ES-type exhibited significantly shorter latency (Fig. 6; unpaired t-test, p < 0.05). In contrast, no significant difference in latency was observed between the two types among SR-responsive neurons (unpaired t-test, p = 0.5244).

**Figure 6.**
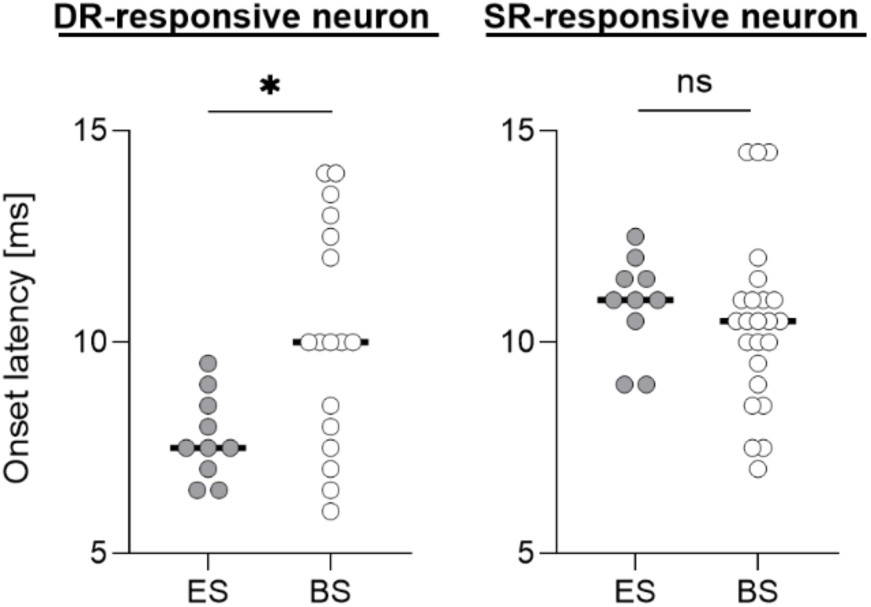
Onset latency of DR- and SR-responsive neurons in M1. Onset latency of extension-suppressed (ES) and bilaterally suppressed (BS) neurons. Left: DR-responsive, right: SR-responsive neuron. Numbers in bars indicate the number of neurons. Dots represent individual neurons. * p<0.05, ns: not significant, unpaired t-test.

Overall, we conclude that the ES-biased distribution is a distinctive feature of DR-responsive neurons during the Static hold period. This pattern may reflect the selective preservation of fast-conducting muscle afferent input from the antagonist (i.e., extensor muscle afferents during wrist flexion), potentially to enable rapid adjustments to fluctuations in motor output during static holding and to support precise control of wrist position or force output.

## Discussion

Our study demonstrates that sensory input from muscle afferents can elicit reliable neuronal activation in M1, as evidenced by LFPs and spiking responses evoked by muscle nerve stimulation. Importantly, this afferent-driven activation was significantly attenuated during voluntary movement, providing direct evidence of sensory gating of muscle afferent input at the level of M1—an effect previously suggested but not clearly shown. Interestingly, a small population of M1 neurons retained their responsiveness in a context-dependent manner. In particular, responsiveness to inputs from antagonist muscles remained effective during the Static hold epoch of the task, when fine control of joint angle was necessary to maintain posture. This finding implies that selected afferent signals may be preserved to support postural stabilization.

Taken together, our results suggest that M1 engages in multisensory gating, modulating both cutaneous and muscle afferent inputs to suppress reafferent signals during movement. The selective preservation of antagonist muscle input during the Static hold period points to a task-specific modulation of gating, which may contribute to the precise control of static hand posture.

### Characteristics and significance of muscle afferent inputs to M1

In the present study, we observed LFPs in M1 evoked by electrical stimulation of the DR nerve, which innervates muscle spindles and Golgi tendon organs of extensor muscles. These afferents convey information about muscle mechanical states—such as position and force—collectively referred to as the “kinesthesia” (Bastian, 1887; McCloskey, 1981). Our findings suggest that kinesthetic feedback from the periphery reaches M1 in awake, behaving non-human primates.

Notably, the latencies of DR-evoked LFPs were significantly shorter than those of SR-evoked LFPs when stimulation was applied at equivalent intensities (Fig. 1C). This latency difference, approximately 1 ms, is consistent with prior observations at the spinal level (Confais et al., 2017), suggesting a peripheral or spinal origin. Given the small latency difference, one might assume that cutaneous and muscle afferents ascend to M1 via a common supraspinal pathway. However, this interpretation should be made with caution, as similar conduction times do not necessarily imply shared anatomical routes. Conduction delays may still differ depending on factors such as the number of synaptic relays or the conduction velocity of individual axons within each pathway.

Crucially, we found that DR stimulation elicited spiking responses in individual M1 neurons (Fig. 4A), indicating that muscle afferent input can reach M1 with sufficient strength to drive postsynaptic firing under awake, behaving conditions. Previous studies have shown that M1 neurons respond to mechanical perturbations involving both muscle and cutaneous stimulation (Cheney & Fetz, 1984; Conrad et al., 1975; Evarts & Tanji, 1976), and our results align with these findings. Moreover, we identified distinct populations of neurons that responded selectively to either muscle or cutaneous afferent inputs, supporting the notion that M1 integrates signals across multiple somatosensory modalities to support sensorimotor transformation.

### Processing and Gating of Muscle Afferent Signals in M1

We found that DR-evoked LFPs in M1 were significantly suppressed during voluntary movement compared to Rest. This finding, together with our previous reports showing suppression of SR-evoked LFPs during movement (Figs. 3A, B, right; see also Seki & Fetz, 2012), confirms that both muscle and cutaneous afferent signals represented in M1 are attenuated during active motor execution.

Both DR- and SR-evoked signals are transmitted to M1 via the dorsal column–medial lemniscus pathway. Given that both pathways share key relay stations—including the cuneate nucleus (CN), thalamus, and primary somatosensory cortex (S1)—these regions are likely loci of the observed suppression. For cutaneous inputs, suppression has been reported at both S1 and CN, with SR-evoked LFPs reduced by ∼60% during movement (Kubota et al., 2024; Seki & Fetz, 2012). In M1, the same signal was reduced by ∼77% in our study. Although the data were collected from different animals, the use of identical behavioral tasks allows for cautious comparison. In light of this, the stronger suppression in M1 suggests additional attenuation beyond CN and S1, potentially at the S1–M1 corticocortical projection or via direct thalamocortical inputs that bypass S1(Asanuma & Mackel, 1989).

For muscle afferent inputs, the exact site of suppression remains unclear, as suppression in CN or S1 has not been reported. However, based on our previous recordings from the cervical spinal cord (Confais et al., 2017), it is plausible that suppression occurs at supraspinal levels. Interestingly, some spinal interneurons even show facilitation of muscle afferent input during voluntary movement (Confais et al., 2017; Tomatsu et al., 2023), whereas cutaneous input is generally suppressed. If muscle afferent signals are indeed facilitated at the spinal level, the robust reduction observed in M1 implies that additional gating must occur upstream, possibly at CN or higher centers. Future work targeting DR-evoked responses in CN and S1 will help clarify this issue.

### Modulation of individual neuronal sensitivity to muscle afferent input in M1

While LFPs primarily reflect synaptic inputs, spiking activity represents the output of a given region (Logothetis et al., 2001). Our finding that DR stimulation evoked spikes in M1 neurons indicates that muscle afferents can drive M1 output under normal behavioral conditions. While electrical nerve stimulation synchronously activates a large population of afferent fibers and may strongly drive cortical neurons, prior studies using naturalistic perturbations have shown that even asynchronous, weaker afferent activity—generated by self-initiated or externally applied movements—can evoke spiking in M1 neurons (Fetz et al., 1980; Flament & Hore, 1988; Lemon et al., 1976). These findings suggest that the M1 responses observed in our study are not simply artifacts of artificial stimulation but reflect physiologically relevant sensory-motor processing. From this perspective, sensory gating in M1 may serve to suppress destabilizing input and preserve task-relevant sensorimotor coordination.

Interestingly, we found that evoked responses in M1 were more strongly preserved during the Static hold epoch than during Active movement. It has been proposed that different feedback loops support distinct movement modes: trans-spinal loops dominate during dynamic actions, while trans-cortical loops are preferentially engaged during static holding (Oya et al., 2020). Accordingly, preserved responses during the Static hold epoch may contribute to trans-cortical feedback, while gating of the remaining majority of inputs may still prevent destabilizing influences from unexpected inputs.

### Differential role of sensory gating in M1 between ballistic and static movements

During the Active movement epoch of the task, which involved ballistic wrist movements, the majority of M1 neurons responsive to muscle afferent input showed marked suppression of evoked activity in both flexion and extension trials (Fig. 5C; “BS” pattern). This indicates that sensory input from both agonist and antagonist muscles is uniformly gated in M1 during rapid, goal-directed movement. Given the pronounced differences in the connectivity patterns and reflex functions of afferent inputs from agonist versus antagonist muscles—for example, their distinct roles in autogenic excitation versus reciprocal inhibition (Gottlieb et al., 1982; Mendell & Henneman, 1971)—the observed uniform suppression of both inputs in M1 during ballistic movement is somewhat surprising and counterintuitive. However, it supports the notion that M1 output during ballistic movements relies primarily on descending motor commands, rather than afferent-driven activation (Evarts & Fromm, 1977), likely due to the temporal constraints imposed by sensory feedback delays (Wolpert & Flanagan, 2001). This is consistent with the idea that trans-cortical feedback loops are less influential than trans-spinal pathways during fast movements (Oya et al., 2020).

In contrast, during the Static hold phase of the task, when monkeys were required to maintain wrist position with high precision, a subset of M1 neurons preserved their evoked responses selectively during extension trials. Importantly, these preserved responses were driven by fast-conducting afferent input from antagonist muscles (Fig. 5C; “ES” pattern). This selective preservation aligns with previous studies showing that specific M1 neurons encode mechanical loads primarily during posture maintenance (Kurtzer et al., 2005). Additionally, β-band cortico-muscular coherence increases during static holding (Baker et al., 1997), and trans-cortical feedback is more dominant under such conditions (Oya et al., 2020), both supporting a role for preserved cortical responsiveness during static motor output.

Interestingly, previous work in the cervical spinal cord has shown that presynaptic inhibition of DR afferent input varies depending on whether the signal originates from agonist or antagonist muscles. In particular, inputs from antagonist muscles are suppressed at the spinal level, contrasting with their preserved influence on M1 responses during static holding in the present study. This dissociation suggests that muscle afferent input from antagonist muscles may serve different functions at spinal versus cortical levels. For instance, muscle spindle activation in antagonistic muscles—triggered by passive stretch during posture—may convey fine-grained information about muscle length that is selectively routed to M1 to support long-latency reflexes and precise postural control (Shemmell et al., 2010). Such feedback may be too slow to be effectively integrated at the spinal level, where rapid, autogenic responses predominate (Pruszynski & Scott, 2012). Accordingly, spinal circuits may suppress these inputs, whereas cortical pathways preserve them to subserve context-dependent control of hand posture.

## Supporting information

supplementary informations

## Acknowledgements and Funding

We thank K. Oida for animal care and technical assistance on the surgery. This work was supported by JST SPRING (grant numbers JPMJSP2116 [to J.Y.]), and by a grant-in-aid from the Japan Society for the Promotion of Science (JSPS) (grant numbers 26120003, 23H05488, 24K21313 [to K. S.], 22H03500, 23H04372 [to S. Kubota], 21K15626 [to S. Kikuta]). The authors used ChatGPT (OpenAI) to assist with language editing and grammar correction during manuscript preparation.

## Author contributions

J. Y., S. Kubota., and K. S. designed research; J. Y., S. Kubota., S. Kikuta and W. H. performed research; J. Y. analyzed data; J. Y., S. Kubota and K. S. wrote the paper.

