## supplementary informations for "Dynamic and context-dependent modulation of proprioceptive input in primate primary motor cortex"

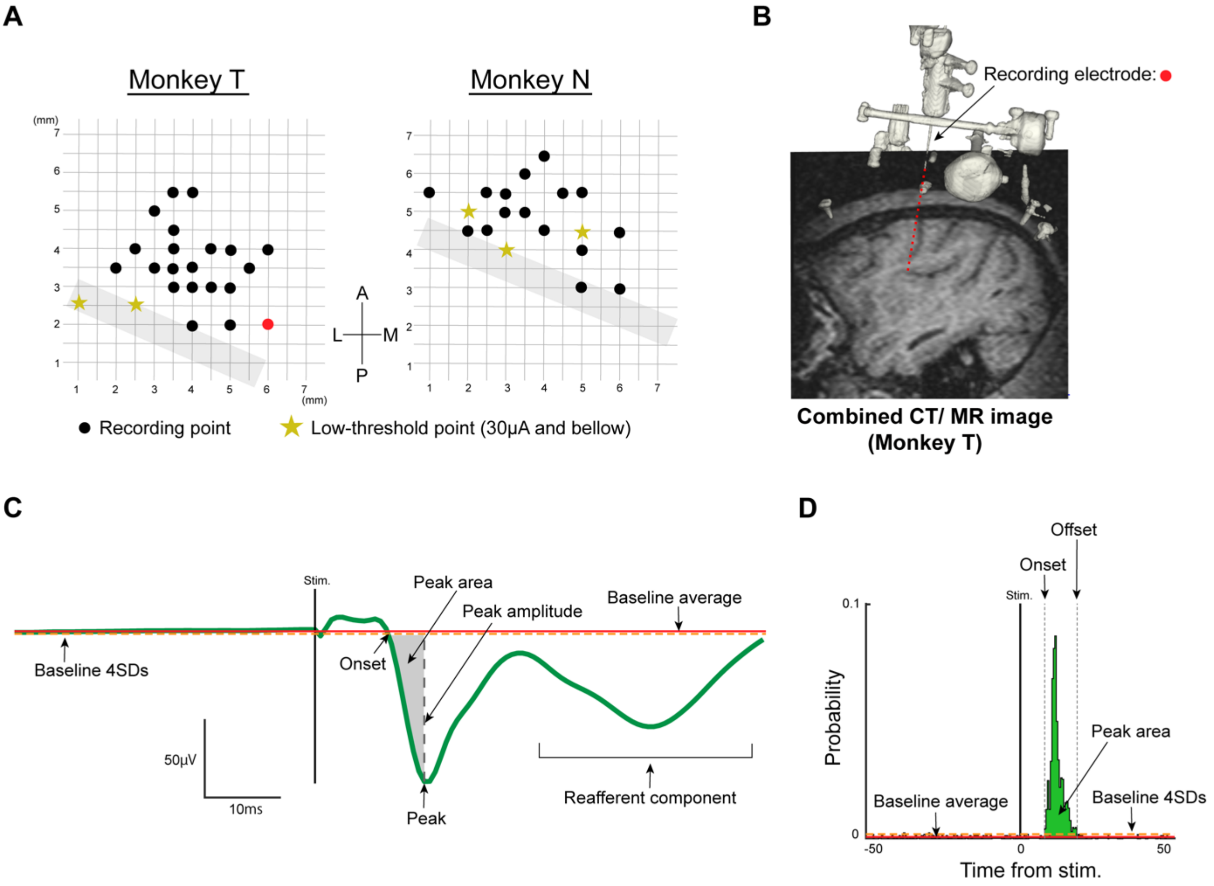


**Figure S1. Recording procedures**

**A,** X-Y coordinates of each electrode-insertion point within the recording chamber in monkeys T and N. Red dot indicates the site shown in **B.** Grey bars denote estimated central sulcus location. A, anterior; P, posterior; L, lateral; M, medial.

**B,** Sagittal view of electrode trajectory. MR image aligned with electrode trajectory (black arrow), reconstructed from CT data. The red dotted line shows the electrode track.

**C,** Peak area measurement for evoked LFPs. The size of the evoked potentials (grey shading) was measured as the peak area under the baseline from the onset to the waveform peak. Example shown for averaged DR-evoked LFP.

**D,** Peak area measurement in PSTH. PSTH peak area (green shading) calculated from the area above the baseline average (red line) from onset to offset. Example shown for DR-responsive neuron.

**Table S1. Suppression patterns of evoked responses in M1 neurons during wrist flexion and extension movement**

| | DR-responsive neuron | | SR-responsive neuron | |
|---|---|---|---|---|
| | **AM** | **SH** | **AM** | **SH** |
| No modulation | 1 (1/0) | 2 (2/0) | 2 (0/2) | 5 (2/3) |
| Flexion-suppressed | 0 (0/0) | 1 (1/0) | 2 (1/1) | 4 (3/1) |
| Extension-suppressed | 3 (3/0) | 10 (4/6) | 5 (4/1) | 10 (7/3) |
| Bilaterally suppressed | 26 (11/15) | 17 (8/9) | 34 (18/16) | 24 (11/13) |

Each row shows the number of classified M1 neurons in the Active movement and Static hold epochs (SR, n = 43; DR, n = 30). Numbers in parentheses indicate the number of neurons from monkeys T and N, respectively. AM, Active movement epoch; SH, Static hold epoch.

**Table S2. Differences in the proportions of flexion- and extension-suppression neurons responsive to DR nerve stimulation during the Static hold epoch**

| Type | DR-responsive neuron | | p-value |
|---|---|---|---|
| | Directional | Non-directional | |
| Flexion-suppressed | 1 | 29 | <0.01# |
| Extension-suppressed | 10 | 20 | |

Each row shows a contingency table of the suppression types of DR-responsive neurons during the Static hold period (n = 30). #, Fisher's exact test.

**Table S3. Differences in the proportions of flexion- and extension-suppression neurons responsive to SR nerve stimulation during the Static hold epoch**

| **Type** | **SR-responsive neuron** | | **p-value** |
|---|---|---|---|
| | **Directional** | **Non-directional** | |
| Flexion-suppressed | 4 | 39 | 0.1433# |
| Extension-suppressed | 10 | 34 | |
